# Adverse neurological effects of short-term sleep deprivation in aging mice are prevented by SS31 peptide

**DOI:** 10.1101/2020.06.04.130435

**Authors:** Jinzi Wu, Yan Dou, Warren C. Ladiges

## Abstract

Sleep deprivation is a potent stress factor that disrupts regulatory pathways in the brain resulting in cognitive dysfunction and increased risk of neurodegenerative disease with increasing age. Prevention of the adverse effects of sleep deprivation could be beneficial in older individuals by restoring healthy brain function. We report here on the ability of SS31, a mitochondrial specific peptide, to attenuate the negative neurological effects of short-term sleep deprivation in aging mice. C57BL/6 female mice, 20 months old, were subcutaneously injected with SS31 (3mg/kg) or saline daily for 4 days. Sleep deprivation was 4 hours daily for the last 2 days of SS31 treatment. Mice were immediately tested for learning ability followed by collection of brain and other tissues. In sleep deprived mice treated with SS31, learning impairment was prevented, brain mitochondrial ATP levels and synaptic plasticity regulatory proteins were restored, and ROS and inflammatory cytokines levels were decreased in the hippocampus. The observations suggest possible therapeutic benefits of SS31 for alleviating adverse neurological effects of acute sleep loss.

## Introduction

Studies have shown that sleep plays a vital role in brain plasticity and memory [1,2] and ensures efficacy of electrical firing within the neuronal synapse [3,4]. Disruption of normal sleep patterns triggers inflammatory pathways and interferes with synaptic transmission in the hippocampus, a specific area of the brain crucial for encoding and memory storage [5,6]. Sleep deprivation has been shown to induce mitochondrial dysfunction that leads to increased reactive oxygen species (ROS) and decreased ATP production [7]. The neurological damage induced by sleep deprivation is especially prevalent in the elderly due to increased incidence of sleep disorders and exaggerated neuroinflammatory responses [8–12]. As a result, finding a rational treatment for sleep deprivation could be highly beneficial by decreasing the risk for neurodegenerative conditions such as Alzheimer’s disease.

SS31 (D-Arg-dimethylTyr-Lys-Phe-NH2), a synthetic mitochondrial-specific peptide, has been shown to improve synaptic and cognitive impairments in mice and rats exposed to lipopolysaccharide and isoflurane. The mechanism was mainly through reducing neuroinflammation and oxidative stress in the hippocampus, protecting the hippocampal neuron mitochondria, and enhancing synaptic plasticity and its related signaling pathways [13–15]. In another study, aged mice treated with SS31 displayed significantly improved neurovascular coupling responses and cognitive functions, including spatial working memory and motor skill learning [16]. Effect of SS31 on learning impairment associated with short-term sleep deprivation in aging mice has not yet been investigated. We show in this report that SS31 prevented learning impairment in old mice after short-term sleep deprivation, and provide data suggesting the protective effects are mediated by preserving mitochondrial integrity and synaptic function.

## Results & Discussion

### Sleep-deprived learning impairment was prevented in aging mice treated with SS31

In order to determine if SS31 could prevent learning deficits induced by short-term sleep deprivation (SD), 20-month-old mice were treated with SS31 (3 mg/kg) by subcutaneous injection once daily for 4 days. On the third and fourth days of treatment, mice were sleep deprived for 4 hours each day. Immediately following the last sleep deprivation session, a spatial navigation task, designated as the Box maze [17,18] was used to measure learning impairment, with escape time as the readout. Non-SD mice showed robust learning ability as indicated by the decreased escape times starting in trial 2 compared to SD mice (Figure. 1). When SD mice were treated with SS31, learning impairment was significantly alleviated and more closely resembled learning ability in non-SD mice. These results suggest that SS31 can prevent learning impairment induced by short-term sleep deprivation in aging mice.

**Figure 1.**
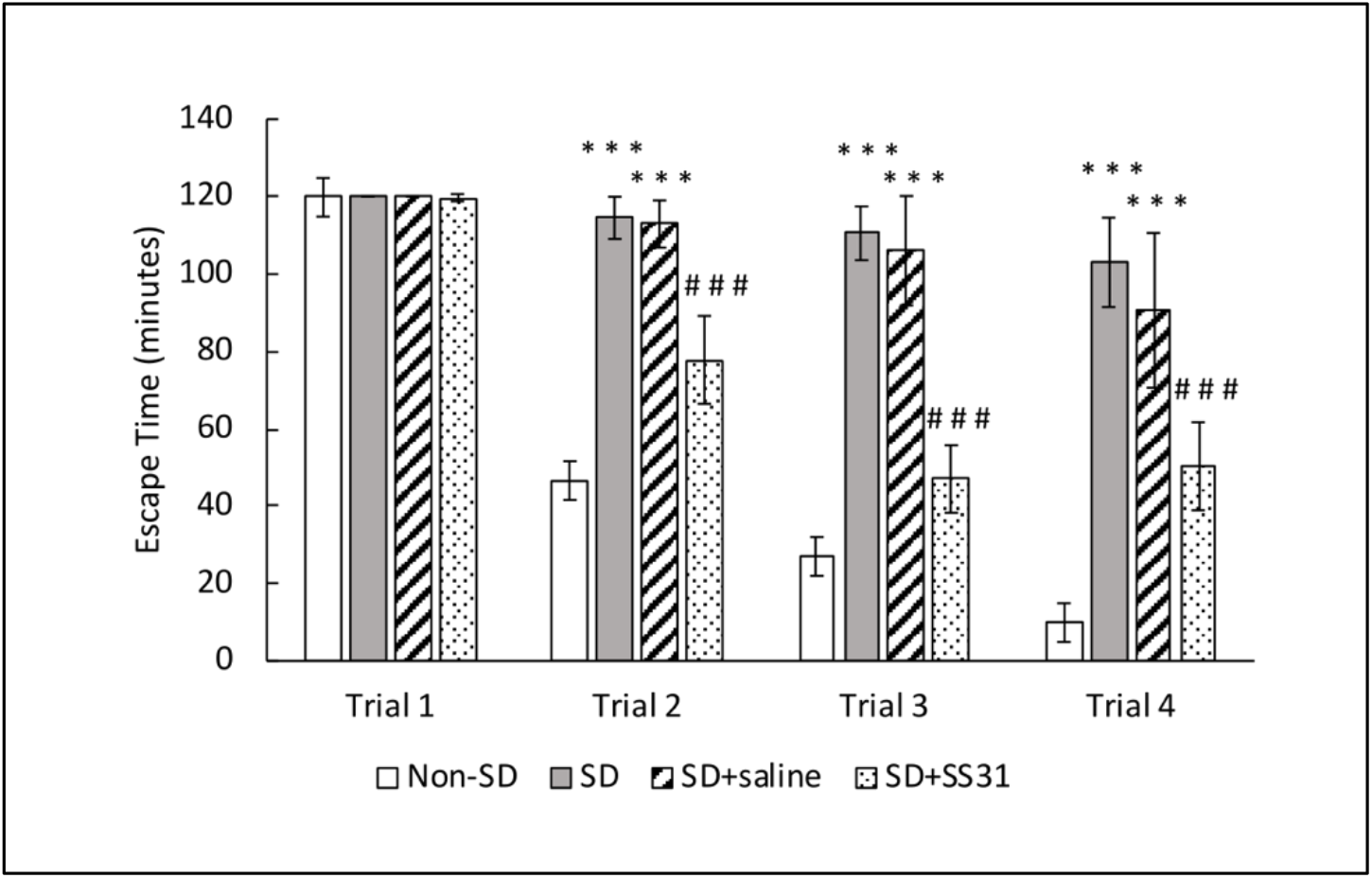
Sleep-deprived learning impairment was prevented in aging mice treated with SS31. Mice were given 4 trials to escape the box maze for a maximum of 2 minutes each, and their escape times were recorded. The data represent the mean ± SEM (n = 10 per group). ***p < 0.001, statistically significant difference between SD group/SD+Saline group and Non-SD group. ###p < 0.001, statistically significant difference between SD+SS31 group and SD group/SD+Saline group.

### ATP was increased and ROS decreased in mitochondria from brain and liver of sleep-deprived mice treated with SS31

Sleep-deprived (SD) mice had a significant decrease in ATP synthesis compared to non-SD mice in the brain (Figure. 2A), suggesting disruption of mitochondrial function and inefficient energy production. On the other hand, ATP levels were restored in SD mice treated with SS31 to levels similar to non-SD mice. In order to determine if there was an increase in ROS that might be related, we measured ROS production in the brain and showed that SD mice had significantly higher levels of mitochondrial ROS in brain compared to non-SD mice (Figure. 2B), suggesting a correlation with disruption of ATP production. SD mice treated with SS31 showed a significant decrease in ROS, with levels similar to non-SD mice, further indicating the disruptive sensitivity of mitochondria to sleep deprivation. We also interrogated the liver for the above molecules in order to determine if mitochondrial function was affected by SD in nonneuronal systemic tissues that might indirectly impact neuronal function. Results were similar to that seen in the brain, with SD causing lower ATP levels and higher levels of ROS (Figure. 2C and D), both of which were rescued by treatment with SS31.

**Figure 2.**
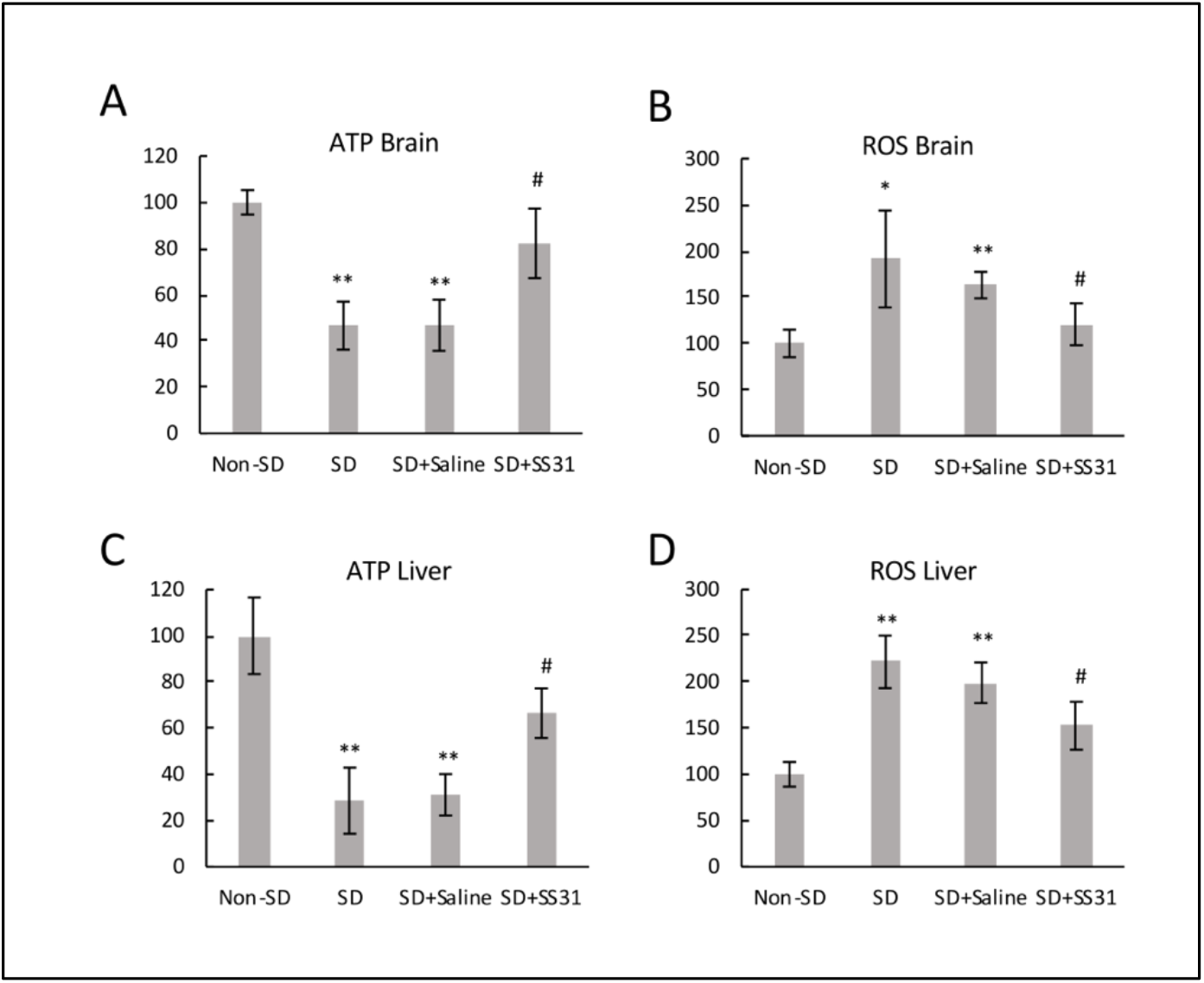
ATP was increased and ROS decreased in mitochondria from brain and liver of sleep-deprived mice treated with SS31. ROS and ATP levels were measured using isolated mitochondria from brain and liver. The y-axis shows percentages, with the non-SD group being 100 percent. **(A)** ATP level in brain tissue. **(B)** ROS level in brain tissue. **(C)** ATP level in the liver tissue and **(D)** ROS level in liver tissue. All data represent the mean ± SEM of three independent triplicate experiments. **p < 0.01, *p < 0.05, statistically significant difference between SD group/SD + Saline group and Non-SD group. #p < 0.05, statistically significant difference between SD+SS31 group and SD group/SD + Saline group.

### Regulatory proteins for synaptic plasticity were restored, and inflammatory cytokines decreased in the hippocampus of sleep-deprived mice treated with SS31

To further investigate the cellular mechanism of how SS31 attenuated learning impairment induced by sleep deprivation in aging mice, we tested expression levels of three known regulators of synaptic plasticity. N-methyl-D-aspartate (NMDA) receptor is a glutamate receptor that plays a vital role in regulating synaptic plasticity and subsequent function in the hippocampus [4,19]. CREB (cAMP-response element binding), a transcription factor downstream of the cAMP/PKA signaling pathway, and brain-derived neurotrophic factor (BDNF) are other regulators of synaptic plasticity, playing vital roles for learning and memory [3,20,21]. Compared to non-SD mice, SD mice had significantly lower levels of NMDA receptor, p-CREB, and BDNF (Figure. 3A, B, C), indicating the negative effects of short-term sleep deprivation on synaptic plasticity-related regulation that might be the result of high sensitivity to ROS-mediated inflammation in the brain. Treatment with SS31 greatly attenuated the decrease in plasticity regulator protein expression induced by sleep deprivation, suggesting an important role of mitochondrial function in synaptic plasticity.

**Figure 3.**
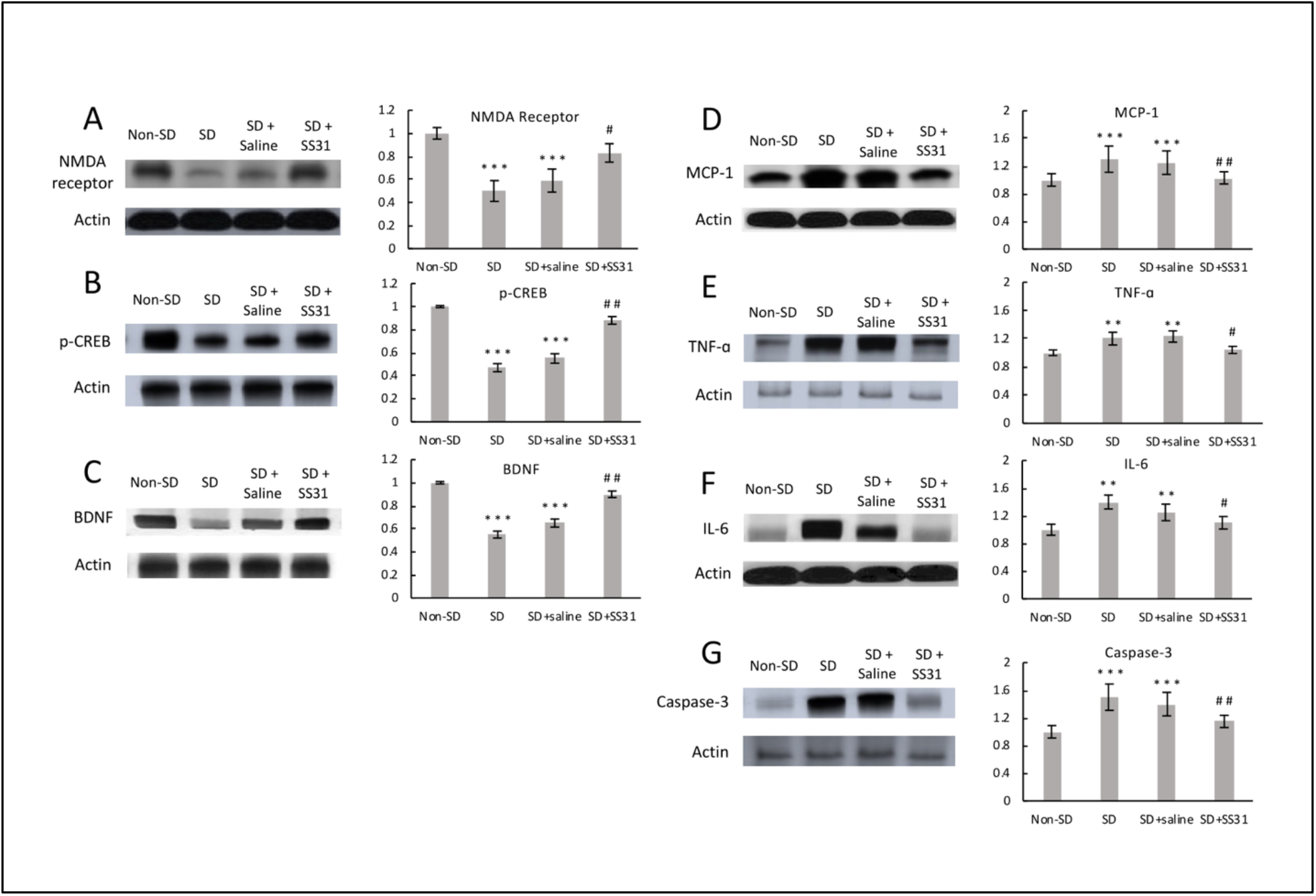
Regulatory proteins for synaptic plasticity were restored, and inflammatory cytokines decreased in the hippocampus of sleep-deprived mice treated with SS31. Western blotting was performed on hippocampus using specific antibodies for the detection of **(A)** NMDA receptor, **(B)** p-CREB, **(C)** BDNF, **(D)** MCP-1, **(E)** TNF-α, **(F)** IL-6, and **(G)** Caspase-3, with Actin used as loading control. The densitometry values for the proteins were normalized to those of Actin, with non-SD group being 1. All data represent the mean ± SEM (n=5 per group). ***p < 0.001, **p < 0.01, statistically significant difference between SD group/ SD+Saline group and Non-SD group. ##p < 0.01, #p < 0.05, statistically significant difference between SD+SS31 group and SD group/ SD+Saline group. Blots have been cropped and originals are available.

Neuroinflammation caused by sleep deprivation is associated with synaptic impairment, especially in the hippocampal area which is particularly sensitive to its detrimental effects [6,22]. To examine the effects of SS31 on inflammation in the hippocampus, we measured levels of three inflammatory cytokines-MCP-1, TNF-α, and IL-6. Compared to non-SD mice, SD mice had significantly higher levels of MCP-1, TNF-α, and IL-6 expression (Figure. 3D, E, F), which might be the result of increased ROS in neurons. Cytokines such as TNF-α and IL-6 have been shown to induce changes in hippocampal dependent learning and memory tasks [23]. Increased hippocampal inflammation in relation to sleep deprivation has been well studied, and the effect of neuroinflammation on cognitive function can be detrimental [6,13,24]. SD mice treated with SS31 had a significant decrease in hippocampal inflammation, to a level similar to non-SD mice, suggesting that mitochondrial dysfunction appears to be related to the increased inflammation caused by short-term sleep deprivation in aging mice.

Increased presence of ROS could overwhelm antioxidant capacity, resulting in cellular damage [25] and activation of the apoptotic pathway [26]. Because an increase in inflammation can trigger an apoptotic response, we tested for Caspase-3, an enzyme known to be involved in the apoptotic pathway [27]. Mice that were sleep deprived had significantly elevated levels of Caspase-3 expression in the hippocampus compared to non-SD mice (Figure. 3G), indicating activation of the apoptotic neuronal cell death pathway in response to compromised neuronal function caused by short-term sleep deprivation. Caspase-3 was greatly reduced in SD mice treated with SS31, suggesting a relationship between sleep deprivation-induced mitochondrial dysfunction, neuroinflammation and elimination of compromised neurons.

Although we do not have data to confirm a mechanistic sequence of events, the sleep deprivation-induced increase in ROS related to mitochondrial dysfunction most likely started a molecular cascade of neuronal dysfunction, leading to learning impairment. SS31 is a free radical scavenger with an ability to interact with the inner mitochondrial membrane to modulate electron flux, increase ATP generation and decrease ROS production [28,29]. Since treatment with SS31 reversed the detrimental effect of sleep deprivation in aging mice, it is logical to suggest a cause and effect relationship between sleep deprivation-induced mitochondrial dysfunction, oxidative stress and learning impairment.

In summary, our studies have linked sleep deprivation with SS31 and aging. The data suggest that short-term sleep deprivation is associated with significant adverse changes in cognitive function, mitochondrial ROS production and energy output, regulation of synaptic plasticity, and inflammatory pathways against an aging background. The study sheds light on the possible acute therapeutic benefits of SS31 for sleep loss conditions and related cognitive decline with increasing age, because of the peptide’s ability to preserve mitochondrial integrity in vital synaptic areas.

## Material and Methods

### Animals and treatment schedule

C57BL/6 female mice, 20 months of age, were obtained from the Aging Rodent Colony of the National Institute on Aging managed by Charles River Inc. Animals were maintained in a specific pathogen free housing facility at the University of Washington with standard rodent chow *ad lib*, reverse osmosis water via an automatic watering system, and room temperature maintained at 70-72 degrees F. Mice were allowed to acclimate to the housing conditions for at least two weeks before entering into any experiments. All animal experiments were approved by the University of Washington Institutional Animal Care and Use Committee.

Mice were treated with SS31 (Stealth BioTherapeutics, Newton, MA) by subcutaneous injection at a dose of 3 mg/kg once daily for four days (Supplement Figure 1). On the third and fourth days of treatment, mice were sleep deprived for 4 hours each day. Immediately following the last sleep deprivation session, a spatial navigation task was used to measure learning impairment.

### Sleep deprivation procedure

The sleep deprivation procedure was carried out as previously described [17]. Based on a 12:12 dark/light cycle, the animals were sleep deprived starting 4 hours after the lights came on for a total of 4 hours daily for 2 days. Sleep deprivation was achieved through continual low stress sleep disturbances including cage tapping and gentle stroking of the back with a small brush. The Box maze assay was conducted immediately following the last sleep deprivation session.

### Spatial navigation Box maze

The Box maze spatial navigation task was conducted as previously described [18] to determine learning impairment caused by sleep deprivation. Briefly, the Box maze consists of a rectangular clear hard-plastic box (26.5 cm width, 30.5 cm length and 29.2 cm height). Each side of the box has two holes and each hole has a distinctive decoration placed above it. The holes were placed and centered 3 cm from the bottom of the cage. During the procedure, 7 of the holes were blocked with one escape hole open to a tube leading to an escape cage. Testing consisted of four 120-second trials. A trial was scored as completed when all four paws were inside the escape hole. The time (latency) to complete the trial was then recorded. If the mouse was unable to find the escape hole it was shown the escape hole and given a latency time of 120 seconds. Between trials, odor markers were removed from the maze with 70% ethanol.

### Western blot

Hippocampus was isolated from the mouse brain within 2 minutes of euthanasia, and snap frozen at −80 °C to be used for Western blots. Protein extraction was carried out using T-PER™ Tissue Protein Extraction Reagent (ThermoFisher Scientific, #78510) The blots were developed using ECL immunochemical detection kit (Bio-Rad, Richmond, CA, USA) and densitometric analysis was conducted to quantify the Western blot immunoreactivity with a scanner and ImageQuant software (Amersham Biosciences, USA). The primary antibodies used were N-methyl-D-aspartate (NMDA) receptor (Cell Signaling Technology, 4207), p-CREB (Cell Signaling Technology, 9198), Caspase-3 (Cell Signaling Technology, 9662S), BDNF (Abcam, ab203573), TNF-α (Novus Biologicals, NBP1-19532), IL-6 (Novus Biologicals, NB600-1131), CCL2/MCP1 (Novus Biologicals, NBP-07035) and actin (Cell Signaling Technology, 4967). The secondary antibody used was goat anti-rabbit IgG-HRP (Santa Cruz Biotechnology, Santa Cruz, CA, USA).

### Isolation of mitochondria

#### Brain

Mitochondria isolation from the whole brain was carried out using Percoll gradient centrifugation as previously reported [30] with slight modifications [31,32]. Brains were removed rapidly and homogenized in 15 ml of ice-cold mitochondrial isolation buffer containing 0.32 M sucrose, 1 mM EDTA and 10 mM Tris-HCl, pH 7.1. The homogenate was centrifuged at 1,330 g for 10 min and the supernatant was saved. The pellet was resuspended in half volume (7.5 ml) of the original isolation buffer and centrifuged again under the same conditions. The two supernatants were combined and centrifuged further at 21,200 g for 10 min. The resulting pellet was resuspended in 12% Percoll solution (Fisher Scientific, 45-001-74) prepared in mitochondrial isolation buffer followed by centrifugation at 6,900 g for 10 min. The obtained soft pellet was resuspended in 10 ml of the mitochondrial isolation buffer and centrifuged again at 6,900 g for 10 min. All of the mitochondrial pellets obtained after centrifugation were either used immediately or frozen at −80°C until analysis. The protein concentration was determined by the Pierce BCA assay kit (ThermoFisher Scientific, #23227).

#### Liver

Mitochondria from the liver were isolated according to a previous described method [33] with slight modifications. Liver tissues were homogenized (1g/10 ml isolation buffer) in mitochondrial isolation buffer containing 15 mM MOPS (pH 7.2), 70 mM sucrose, 230 mM mannitol, and 1 mM K^+^-EDTA. The homogenates were centrifuged at 8,000g for 10 min at 4 °C. The resulting supernatant was further centrifuged at 8,000g for 10 min at 4 °C. The resulting pellet containing crude mitochondria was washed with 10 ml of the isolation buffer followed by centrifugation under the same conditions. The obtained mitochondrial pellet was either used immediately or frozen at −80 °C until further use. The protein concentration was determined by the Pierce BCA assay kit (ThermoFisher Scientific, #23227).

### ROS assay

The ROS assay was carried out using mitochondria isolated from liver and brain with the Reactive oxygen species assay kit (MyBioSource, MBS2540517) according to the manufacturer’s protocol manual. DCFH-DA (2,7-Dichlorofluorescein diacetate) is a fluorescent probe without fluorescence that can freely cross the cell membrane and can be hydrolyzed by intracellular esterase to form DCFH (dichlorofluorescin). In the presence of ROS, DCFH is oxidized to DCF (dichlorofluorescein) which is a strong green fluorescent substance that cannot penetrate the cell membrane. In short, 20uM of DCFH-DA working solution was added to the mitochondrial pellet and incubated for 1 hr at 37°C. The mitochondrial suspension was then centrifuged, and the pellets were washed two times with the working reagent from the kit, and resuspended in the working reagent with equal volume and equal protein concentration on a microplate for fluorescence detection. Fluorescence at 500 nm excitation and 525 nm emission was then read on a fluorescence microplate reader (Synergy H1, BioTek, Winooski, VT, USA).

### ATP assay

The ATP assay was carried out using mitochondria isolated from liver and brain with the ATP Colorimetric/Fluorometric Assay Kit (BioVision, K354-100) according to the manufacturer’s protocol manual. In short, 50 ul of ATP assay reaction mix was added to 50ul of mitochondrial suspension on a microplate and incubated at room temperature for 30 min. Fluorescence at 535 nm excitation and 587 nm emission was then read on a fluorescence microplate reader (Synergy H1, BioTek, Winooski, VT, USA).

### Statistical analysis

All the data are presented as mean ± SEM. Statistical comparisons were performed with the independent t-test. The criterion for statistical significance was considered to be p<0.05.

## Acknowledgements

This work was supported by NIA grant R01 AG057381 (Ladiges, PI). SS31 was the generous gift of Stealth BioTherapeutics, Newton, MA. The technical assistance of Juan Wang and John Morton was greatly appreciated.

## Author Contribution

JW and WL planned the experiments with help from YD. JW carried out the experiments with help from YD. YD and WL wrote the manuscript with help from JW.

## Availability of Materials and Information

There are no restrictions on the availability of materials used in this study. All data generated or analyzed are included in this published article.

