## Supplemental Figure 1 for "Adverse neurological effects of short-term sleep deprivation in aging mice are prevented by SS31 peptide"

### Slide 1
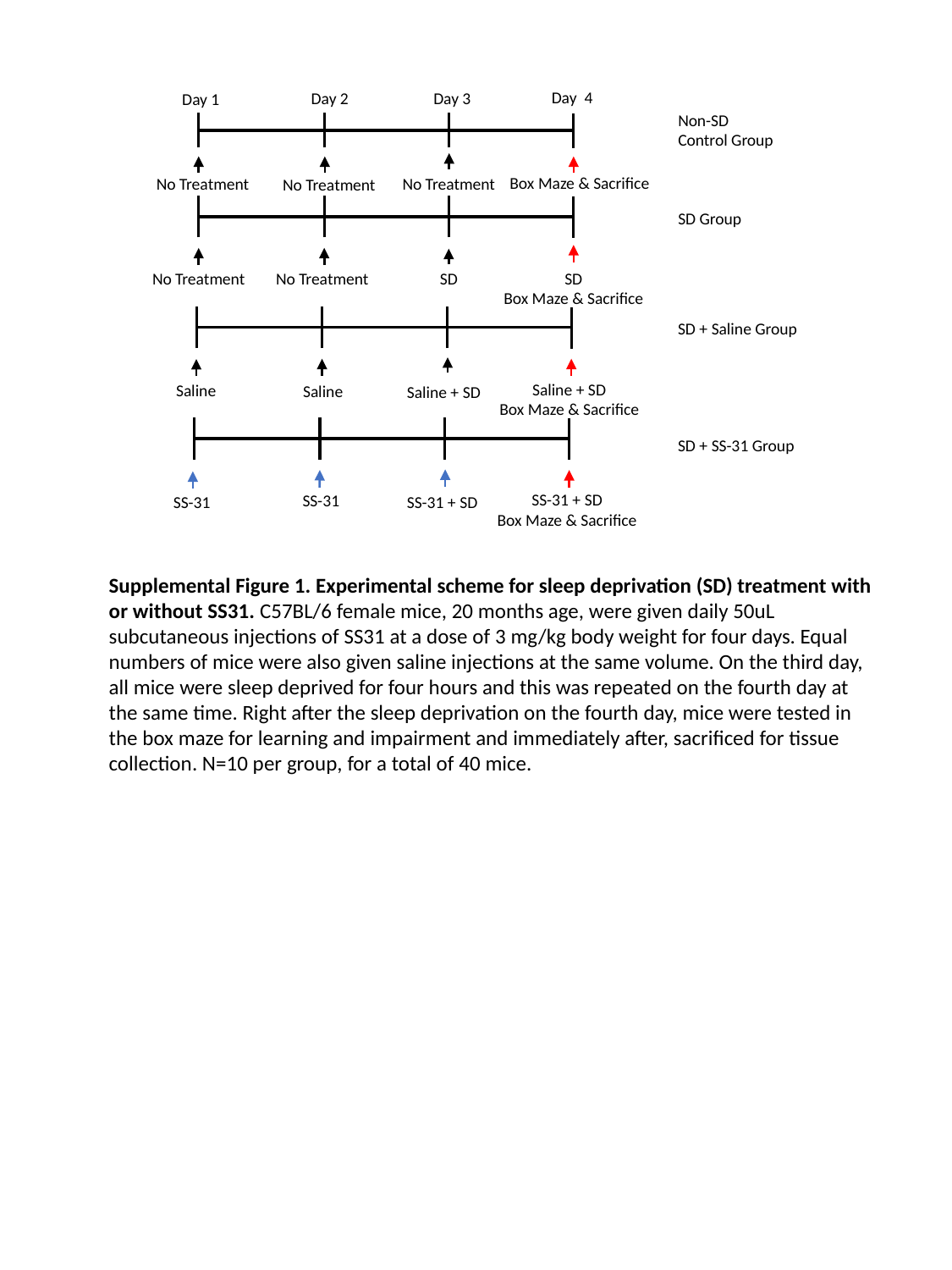

Day 4
Day 2
Day 3
Day 1
Box Maze & Sacrifice
No Treatment
No Treatment
No Treatment
Non-SD
Control Group
SD
Box Maze & Sacrifice
No Treatment
SD
No Treatment
SD Group
Saline + SD
Box Maze & Sacrifice
Saline
Saline
Saline + SD
SD + Saline Group
SS-31 + SD
Box Maze & Sacrifice
SS-31
SS-31
SS-31 + SD
SD + SS-31 Group
Supplemental Figure 1. Experimental scheme for sleep deprivation (SD) treatment with or without SS31. C57BL/6 female mice, 20 months age, were given daily 50uL subcutaneous injections of SS31 at a dose of 3 mg/kg body weight for four days. Equal numbers of mice were also given saline injections at the same volume. On the third day, all mice were sleep deprived for four hours and this was repeated on the fourth day at the same time. Right after the sleep deprivation on the fourth day, mice were tested in the box maze for learning and impairment and immediately after, sacrificed for tissue collection. N=10 per group, for a total of 40 mice.
